# scDock: Streamlining drug discovery targeting cell-cell communication via scRNA-seq analysis and molecular docking

**DOI:** 10.1101/2025.11.20.689638

**Authors:** Chen-Hao Huang, Yen-Jen Oyang, Hsuan-Cheng Huang, Hsueh-Fen Juan

## Abstract

**Summary:** Identifying drugs that target intercellular communication networks represents a promising therapeutic strategy, yet linking single-cell RNA sequencing (scRNA-seq) analysis to structure-based drug screening remains technically challenging and requires substantial bioinformatics expertise. We present scDock, an integrated and user-friendly pipeline that seamlessly connects scRNA-seq data processing, cell–cell communication inference, and molecular docking-based drug discovery. Through a single configuration file, users can execute the complete workflow, from raw scRNA-seq data to ranked drug candidates, without programming skills. scDock automates the identification of disease-relevant ligand–receptor interactions from scRNA-seq data and perfoms structure-based virtual screening against these communication targets using Protein Data Bank (PDB) or AlphaFold-predicted protein structures. The pipeline generates comprehensive outputs at each stage, enabling users to explore intercellular signaling alterations and discover therapeutic compounds targeting specific cell–cell communications. scDock addresses a critical gap by providing an accessible end-to-end solution for communication-targeted drug discovery from single-cell data.

**Availability and Implementation:** scDock is freely available at https://github.com/Andrewneteye4343/scDock. It is implemented in R, Python, shell scripts, and supports Linux systems, including Ubuntu and Debian.

## 1 Introduction

Single-cell RNA sequencing (scRNA-seq) provides high-resolution gene expression profiling at the level of individual cells, offering insights into cellular heterogeneity and complex biological processes that cannot be captured by bulk RNA sequencing (Cao et al., 2017). Recent methodological advances, including cell–cell communication inference (Jin et al., 2025), transcription factor prediction (Aibar et al., 2017), and transcriptional dynamics modeling (Bergen et al., 2020), have further expanded the analytical scope of scRNA-seq, enabling a more comprehensive characterization of transcriptomic landscapes and the mechanisms that underlie diverse biological states. In particular, intercellular communication inference has emerged as a powerful approach for identifying ligand–receptor pairs interactions associated with dysregulated signaling pathways in disease.

Despite offering a broader analytical repertoire than bulk RNA sequencing, scRNA-seq workflows often require complex implementation steps involving multiple programming languages and substantial computational environment setup. These technical demands pose significant challenges to many biologists and can hinder both accessibility and reproducibility. Establishing an intuitive, user-friendly analytical framework is therefore essential to lowering these barriers and improving the usability of scRNA-seq analyses.

Here, we present scDock, a command-line–based integrative toolkit that unifies core scRNA-seq analysis, intercellular communication inference, and drug screening through molecular docking of key signaling molecules. scDock accommodates multiple scRNA-seq input formats and offers accessible options for normalization, dimensionality reduction, clustering, tissue-specific annotation, and data integration. Beyond these core functionalities, the toolkit supports both single-group and comparative analyses of cell–cell communication inference and identifies molecules involved in differential signaling as candidate targets for molecular docking. Targeting critical components of disease-related interactomes has proven to be an effective drug discovery strategy (Lu et al., 2020), and scDock brings this capability into an end-to-end single-cell analysis workflow.

Despite its broad functionality, scDock remains easy to perform analytic workflow and requires only a single configuration file, which provides detailed descriptions and default settings for users with limited computational experience while allowing customization for advanced analyses. Collectively, these features position scDock as a versatile and accessible platform that streamlines single-cell transcriptomic analysis and accelerates the identification of potential therapeutic targets.

## 2 Implementation

scDock is implemented as an integrated toolkit that streamlines complex scRNA-seq analyses and downstream computational workflows for both human and mouse datasets. Its primary goal is to simplify the execution of scRNA-seq analysis and related bioinformatics tasks, thereby reducing the technical burden for users with limited programming experience. To achieve this, scDock is organized into three interconnected functional modules: core scRNA-seq analysis, intercellular communication inference, and molecular docking, all of which operate together to support a fully automated and customizable end-to-end workflow (Fig. 1). These modules are coordinated through R, Python, and shell scripts, enabling the entire process to be executed with a single command and a user-defined configuration file.

**Fig. 1.**
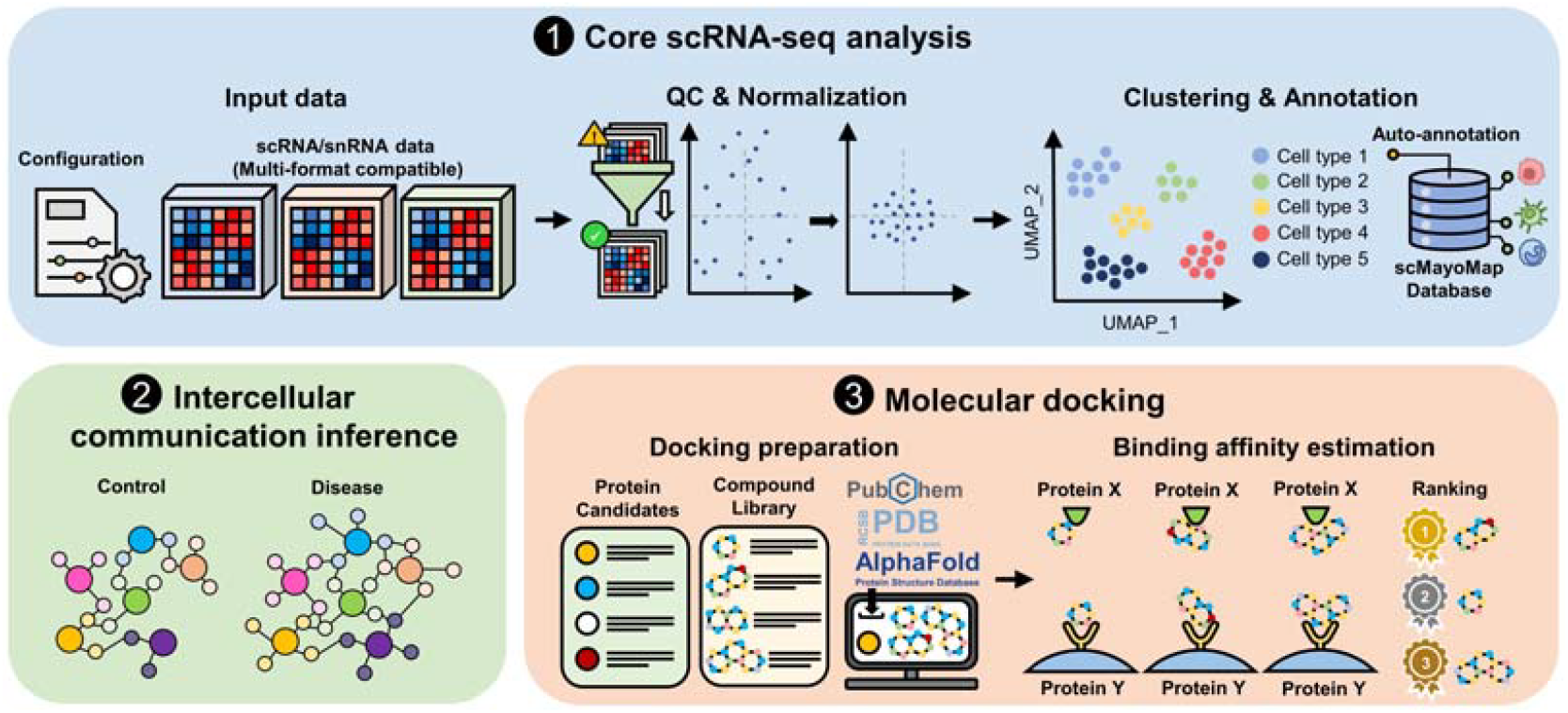
Overview of the scDock analytical workflow. The pipeline consists of three major modules: the core scRNA-seq analysis module, the intercellular communication inference module, and the molecular docking module. The core scRNA-seq analysis module performs quality control, normalization, scaling, dimensionality reduction, clustering, integration, and cell type annotation. The intercellular communication inference module identifies key signaling pathways by comparing signaling probabilities across groups and determines potential therapeutic targets based on the ligand–receptor pairs involved. In the molecular docking module, protein and compound structures are retrieved, preprocessed, and subjected to docking analysis. Compounds are then ranked by their predicted binding affinities to individual target proteins, enabling the identification of candidate drugs that may modulate disease-associated ligand–receptor interactions.

The configuration file specifies all key analytical parameters, including input and output paths, selected analysis methods, algorithms, and the definition of cell types or groups of interest. It provides both default and fully customizable settings, accommodating the needs of novice users while offering flexibility for advanced users. scDock can be downloaded, installed, and executed locally by following the instructions provided on GitHub. Once a scRNA-seq dataset is supplied, the pipeline automatically performs marker gene identification, cell annotation, intercellular communication inference, and molecular docking, producing outputs such as cell-annotated dimensionality reduction plot, summaries of signaling pathways, protein and compound structures, docking results, and other associated files. All intermediate data generated throughout the workflow are also saved automatically, allowing users to reuse, inspect, or extend these results for additional downstream analyses.

### 2.1 Core scRNA-seq analysis

In the core scRNA-seq analysis module, data are processed using the Seurat v5 framework (Hao et al., 2024), which performs quality control, normalization, scaling, dimensionality reduction, clustering, and integration steps when required. To ensure broad compatibility, scDock supports multiple commonly used scRNA-seq matrix formats, including 10x Genomics Cell Ranger outputs (barcodes.tsv.gz, features.tsv.gz, matrix.mtx.gz), hierarchical data format files (.h5), and plain text files (.txt). Both gene symbols and Ensembl identifiers are accepted as input formats. Since non–gene symbol identifiers can reduce annotation accuracy and slow downstream analyses, scDock automatically converts Ensembl identifiers to their corresponding gene symbols during preprocessing. For datasets involving multiple experimental or biological conditions, users may provide a metadata file specifying sample group information. These grouping labels are automatically incorporated into a standardized field named *sample_group*, which can be used directly in downstream comparative analyses and integration workflows.

During clustering and dimensionality reduction, the number of principal components (PCs) selected has a critical impact on the amount of biologically informative variation preserved in the dataset (Stuart et al., 2019, Becht et al., 2019). To determine the optimal number of PCs, scDock uses the geometric elbow method as its default strategy. This approach identifies the point with the greatest perpendicular distance from the line connecting the first and last PCs on the variance-explained plot, as previously described (Zhuang et al., 2022). Because batch effects can introduce unwanted technical variability that obscures true biological signals (Hie et al., 2019), scDock incorporates several integration methods to mitigate such artifacts, including canonical correlation analysis (CCA) (Butler et al., 2018), reciprocal PCA (RPCA) (Hao et al., 2024), and the Harmony algorithm (Korsunsky et al., 2019).

Manual cell type annotation is often labor-intensive, requires substantial domain expertise, and must account for tissue-specific context. To address these challenges, scDock integrates scMayoMap (Yang et al., 2023) for automated cell-type annotation across a broad range of tissues. Beyond the original reference database, scDock further expands the annotation resources to improve performance in disease-specific settings. These extensions currently include neuroblastoma, an uncommon and non–tissue-specific cancer that can originate in multiple organs, and breast cancer, enabling more accurate identification of malignant cells and subtype-specific populations.

### 2.2 Intercellular communication inference

The intercellular communication inference module uses CellChat v2 (Jin et al., 2025) to evaluate ligand–receptor expression patterns across cell types and infer their potential communication networks, including secreted signaling, extracellular matrix-receptor interactions, and cell–cell contact-mediated communication. Identifying ligand-receptor pairs involved in prominent signaling pathways reveals key intercellular interactions that may represent promising therapeutic targets. By default, scDock operates in a single-group mode and computes the signaling pathways with the highest interaction probabilities across all cell types, recording both incoming and outgoing signals as well as global interaction patterns.

A central feature of scDock is its ability to investigate disease-associated signaling pathways as the initial step toward therapeutic target discovery. When a metadata file specifying experimental or biological groups is provided, scDock performs group-wise comparisons to quantify differences in intercellular communication patterns. Signaling pathways exhibiting the greatest shifts in interaction probabilities are identified, and their corresponding differentially active ligand–receptor pairs are designated as candidate therapeutic targets. These targets are then automatically forwarded to the molecular docking module for downstream evaluation. scDock also supports cell type-specific communication analyses. By adjusting the parameters *Run_CellChat_source_celltype* and *Run_CellChat_target_celltype* in the configuration file, users can restrict the analysis to selected source and target populations, enabling more focused and biologically informed investigations of intercellular signaling events.

### 2.3 Molecular docking

In the molecular docking module, scDock uses AutoDock Vina (Trott and Olson, 2010) to estimate the binding affinities between small molecules and the target proteins identified through the intercellular communication analysis. Constructing compound and protein structure libraries is typically time-consuming and requires technical expertise; scDock streamlines this process by providing three options for compound library generation: (i) a built-in FDA-approved compound library, (ii) a customized library generated from Chemical Abstracts Service (CAS) registry numbers, and (iii) user-supplied compound structures.

For the first option, scDock includes a curated list of FDA-approved small-molecule drugs (version 2 September 2025) within its GitHub repository, allowing users to rapidly assemble a ready-to-use library for systematic screening. For the second option, users may supply CAS identifiers, and scDock automatically retrieves the corresponding compound structures from PubChem (Kim et al., 2025) and converted them into AutoDock Vina-compatible formats using OpenBabel (O’Boyle et al., 2011) and RDKit (Rational Discovery LLC, 2025). For the third option, users may directly provide pre-processed compound structures that meet AutoDock Vina input specifications.

For protein structure determination, scDock provides curated reference lists (*ligand_reference*.*csv* and *receptor_reference*.*csv*) containing the best available protein models identified via UniProt-PDB-Mapper (Riziotis, 2025). These structures are retrieved from the SWISS-MODEL Template Library (Bienert et al., 2017). When no experimentally determined structures are available, scDock automatically obtains predicted models from the AlphaFold Protein Structure Database (Jumper et al., 2021). Users may update the curated lists when improved structures becomes available or supply their own protein models directly.

scDock performs compound screening using a global docking strategy, automatically defining the docking grid box for each protein to ensure comprehensive exploration of potential binding sites. During the docking procedure, scDock records all protein and compound structures along with the resulting predicted binding poses and binding affinities. The toolkit also summarizes and ranks the affinities of all screened compounds for each target protein, enabling efficient identification of high-priority therapeutic candidates.

## 3 Application

To demonstrate the utility of scDock, we analyzed a scRNA-seq dataset (GSE218563) from diabetic mouse kidneys (Liu et al., 2023), which includes samples from 16 mice representing both control and diabetic nephropathy (DN) groups. A complete walkthrough of this analysis is provided in the GitHub repository. Using the group information specified in the metadata, we examined differences in intercellular communication between the control and DN groups.

Using a single configuration file, scDock produced the following results. First, it identified major kidney cell populations, including distal convoluted tubule cells, proximal tubule cells, and mesangial cells (Supplementary Figure 1). In the intercellular communication analysis, scDock visualized global signaling networks and revealed pathways with increased interaction probabilities in the DN group relative to controls (Supplementary File 2). Among the differential signaling pathways, the interaction between peptidyl-prolyl cis-trans isomerase A (PPIA) and basigin (BSG) was highlighted as associated with DN, leading to the selection of these two proteins as targets for molecular docking.

scDock then performed molecular docking on the selected targets using a common library assembled from user-provided CAS registry numbers. The pipeline generated binding poses and calculated binding affinities for each compound–protein pair. Notably, the predicted complex between glimepiride and PPIA was visualized using PyMOL (Schrödinger LLC, 2021) to provide additional structural illustration (Supplementary Figure 3).

## 4 Conclusions

We developed scDock as an integrative and user-friendly toolkit that streamlines the complex analytical steps and computational expertise typically required for scRNA-seq data analysis. By unifying single-cell transcriptomic processing, inter-cellular communication inference, and molecular docking–based compound screening, scDock lowers the technical barrier for researchers and enables efficient identification of disease-associated therapeutic targets. Comprehensive documentation, installation instructions, and tutorials are available on GitHub. We anticipate that scDock will expand the accessibility of scRNA-seq–based analyses and accelerate the discovery of potential therapeutics across a wide range of biological and disease contexts.

## Data availability

The scDock source code is publicly available on GitHub at https://github.com/Andrewneteye4343/scDock. The single-cell RNA-seq dataset used in the demonstration analysis was obtained from NCBI GEO database under accession number GSE218563 (https://www.ncbi.nlm.nih.gov/geo/query/acc.cgi?acc=GSE218563).

## Acknowledgements

We thank Dr. Kai-Pu Chen at Center for Computational and Systems Biology, National Taiwan University, Dr. Ching-Chia Yang, Hong-Shian Yang, Hsin-Yo Yuan, and Hong-Ming Tseng for their valuable and constructive feedbacks.

## Author contributions

**CHH:** data curation, investigation, formal analysis, methodology, validation, visualization, writing original draft. **YJO:** methodology. **HCH:** conceptualization, methodology, writing original draft, supervision, project administration, funding acquisition. **HFJ:** conceptualization, methodology, writing original draft, supervision, project administration, funding acquisition.

