## Supplementary Data for "scDock: Streamlining drug discovery targeting cell-cell communication via scRNA-seq analysis and molecular docking"

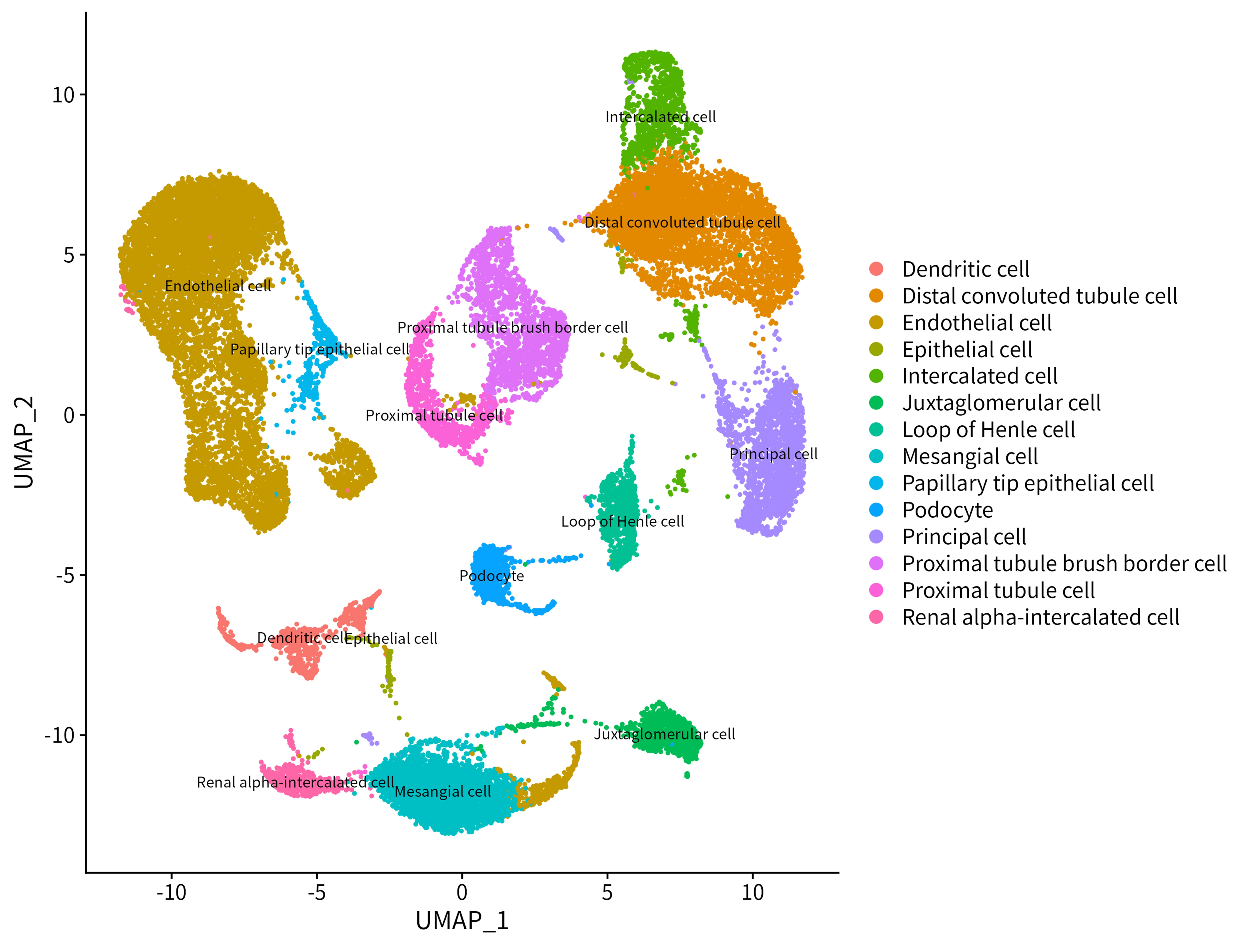

**Supplementary Figure 1. Cell type annotation and distribution of the GSE218563 dataset.**

Each cell type is shown in a distinct color and labeled on the UMAP plot.

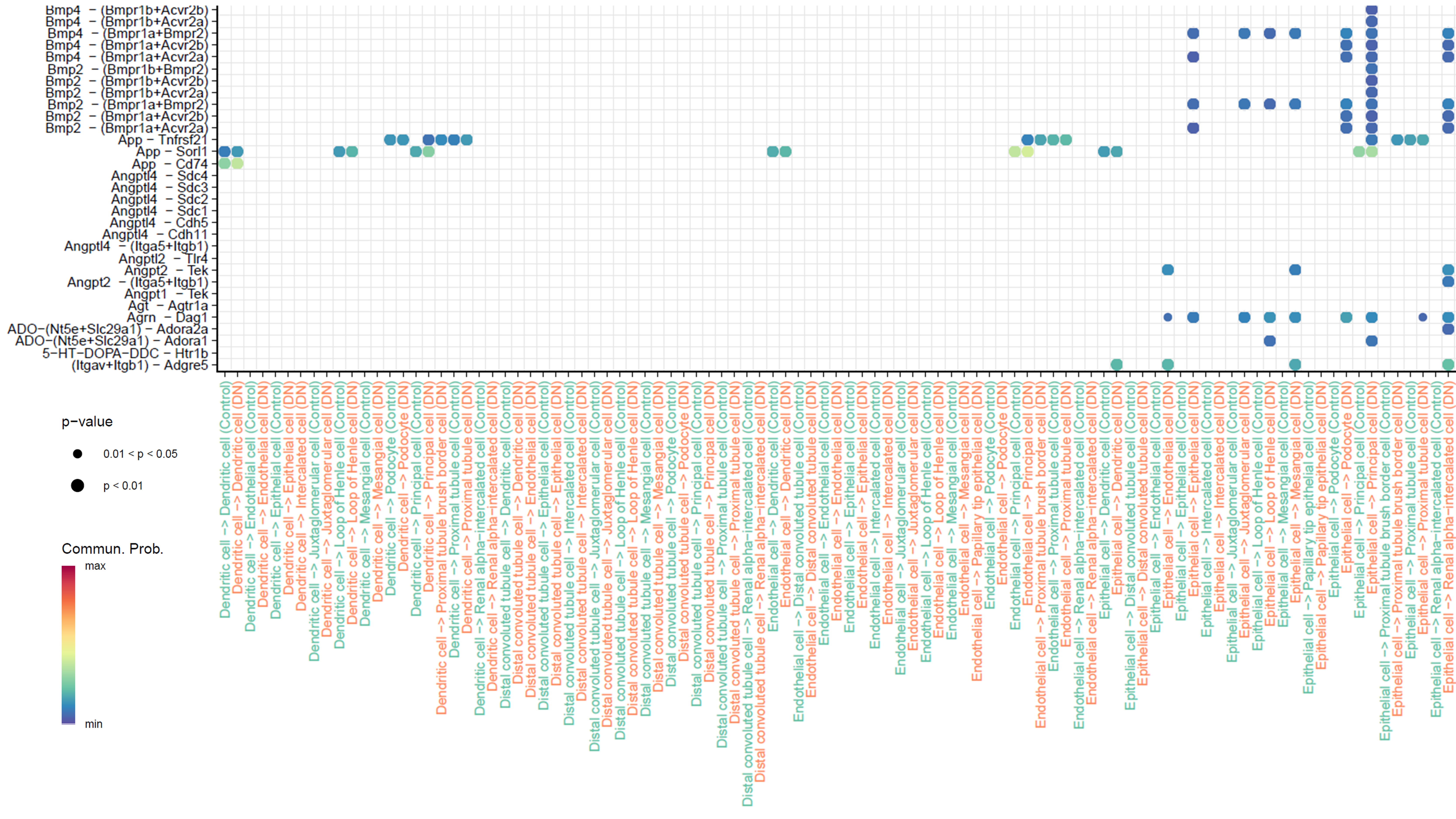

**Supplementary Figure 2. Partial visualization of differential signaling in the diabetic nephropathy (DN) group compared with the control group.**

This bubble plot highlights signaling pathways with higher activity probabilities in the DN group. The control group is shown in green, while the DN group is shown in orange.

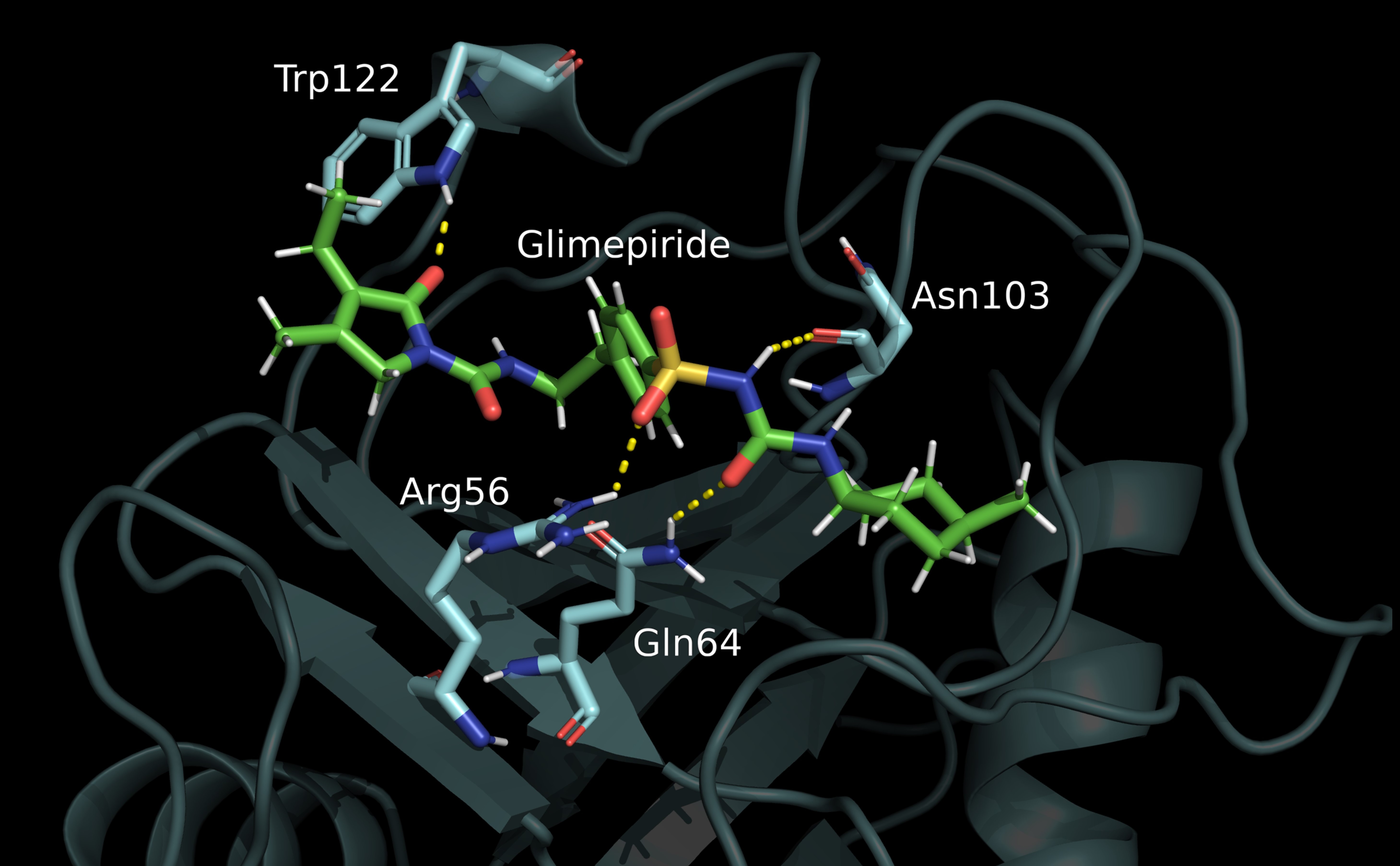

**Supplementary Figure 3. Molecular interaction between glimepiride and the PPIA protein.**

The PPIA protein is shown in cyan, and glimepiride is shown in green. Yellow dashed lines indicate hydrogen bonds between the two molecules.
